# Integrated proteogenomic analysis of metastatic thoracic tumors identifies APOBEC mutagenesis and copy number alterations as drivers of proteogenomic tumor evolution and heterogeneity

**DOI:** 10.1101/301390

**Authors:** Nitin Roper, Shaojian Gao, Tapan K. Maity, A. Rouf Banday, Xu Zhang, Abhilash Venugopalan, Constance M. Cultraro, Rajesh Patidar, Sivasish Sindiri, Alexandr Goncearenco, Anna R. Panchenko, Romi Biswas, Anish Thomas, Arun Rajan, Corey A. Carter, David Kleiner, Stephen Hewitt, Javed Khan, Ludmila Prokunina-Olsson, Udayan Guha

## Abstract

Elucidation of the proteogenomic evolution of metastatic tumors may offer insight into the poor prognosis of patients harboring metastatic disease. We performed whole-exome and transcriptome sequencing, copy number alterations (CNA) and mass spectrometry-based quantitative proteomics of 37 lung adenocarcinoma (LUAD) and thymic carcinoma (TC) metastases obtained by rapid autopsy and found evidence of patient-specific, multi-dimensional heterogeneity. Extreme mutational heterogeneity was evident in a subset of patients whose tumors showed increased APOBEC-signature mutations and expression of *APOBEC3* region transcripts compared to patients with lesser mutational heterogeneity. *TP53* mutation status was associated with APOBEC hypermutators in our cohort and in three independent LUAD datasets. In a thymic carcinoma patient, extreme heterogeneity and increased *APOBEC3AB* expression was associated with a high-risk germline *APOBEC3AB* variant allele. Patients with CNA occurring late in tumor evolution had corresponding changes in gene expression and protein abundance indicating genomic instability as a mechanism of downstream transcriptomic and proteomic heterogeneity between metastases. Across all tumors, proteomic heterogeneity was greater than copy number and transcriptomic heterogeneity. Enrichment of interferon pathways was evident both in the transcriptome and proteome of the tumors enriched for APOBEC mutagenesis despite a heterogeneous immune microenvironment across metastases suggesting a role for the immune microenvironment in the expression of APOBEC transcripts and generation of mutational heterogeneity. The evolving, heterogeneous nature of LUAD and TC, through APOBEC-mutagenesis and CNA illustrate the challenges facing treatment outcomes.

## INTRODUCTION

Metastatic lung adenocarcinoma (LUAD) and thymic carcinoma (TC) have a poor prognosis and treatment options are limited. Understanding the mechanisms by which metastatic LUAD and TC evolve may provide greater insight into tumor progression and may guide novel therapeutic avenues. Previous work characterizing the evolution of primary non-small cell lung cancer (NSCLC) has demonstrated significant intra-tumor heterogeneity^1–3^. However, metastatic lineages are known to occur early in primary tumor development^4^; thus, it is critical to understand the evolution of metastatic NSCLC and other metastatic cancer types. Autopsy programs^5–10^ established to harvest tumor tissue from metastatic sites at the end of life, have demonstrated significant heterogeneity depending on tumor type^11–16^. For example, metastatic pancreatic cancer has high inter-metastatic heterogeneity of genomic rearrangements^12^ but not of driver mutations^17^, whereas metastatic clear cell renal carcinoma has recurrent driver mutations that occur late within individual metastases^18^ resulting in high inter-metastatic heterogeneity. These and other tumor heterogeneity studies have largely focused on whole-exome or genome sequencing approaches. The evolution of tumors at the level of the transcriptome or proteome and the underlying mechanisms that may generate multidimensional heterogeneity remain largely unknown.

Here, we sought to address the following questions: (i) What is the degree of genomic (mutational and copy number), transcriptomic and proteomic heterogeneity within and between metastases of a given patient? (ii) What is the relationship between these three levels of heterogeneity? and (iii) What are the potential drivers of such heterogeneity? To address these questions, we performed rapid (“warm”) autopsies on four patients with lung adenocarcinoma and one patient with thymic carcinoma. The autopsies were initiated within three hours of death, which allowed for procurement of sufficient quantity and quality of DNA, RNA and protein from metastatic tumor tissue for whole-exome and transcriptome sequencing, DNA copy number analysis, and mass spectrometry-based proteomics. Our integrated analysis of the genome, transcriptome and proteome uncovered key mechanisms likely driving proteogenomic heterogeneity within and across metastatic sites.

## RESULTS

### Sampling of metastatic tumors through rapid autopsy protocol

We have established a rapid (“warm”) autopsy protocol for thoracic malignancies (including lung cancers and thymic epithelial tumors, among others) at the NIH Clinical Center. Under this protocol, patients with metastatic disease who are near the end of life receive inpatient hospice care. Upon death, an autopsy is performed within three hours to procure sufficient quantity and high-quality of DNA, RNA and protein from all possible sites of metastatic disease. For this study, we enrolled four patients with lung adenocarcinoma (LUAD) - patients RA000 and RA004 who were previous smokers known to harbor oncogenic *KRAS* mutations and patients RA003 and RA005 who were both non-smokers with *EGFR* mutations. We additionally enrolled squamous cell thymic carcinoma patient RA006, a non-smoker, who had an aggressive disease course marked by no response to treatment and who died within 1.5 years of diagnosis (Supplementary Table 1). All of our study patients were initially diagnosed with stage IV disease, had received chemotherapy and/or targeted therapy (range 2–8 lines of therapy) and were previously enrolled in a clinical trial at the NIH Clinical Center, except for patient RA000 (Supplementary Table 1). For each patient, we harvested between 44–183 metastatic tumor lesions from multiple organs, including lung, liver and kidney (Supplementary Table 1). We selected a total of 40 tumor samples for further analyses based on tumor content by histology (Fig. 1 and Supplementary Fig. 1). Three samples were removed from the study after sequencing and further downstream analysis due to low tumor content. Additional tumor tissue collected at the time of diagnosis, likely from the primary site, was available for whole-exome sequencing in three patients (RA000, RA003 and RA006).

**Figure 1:**
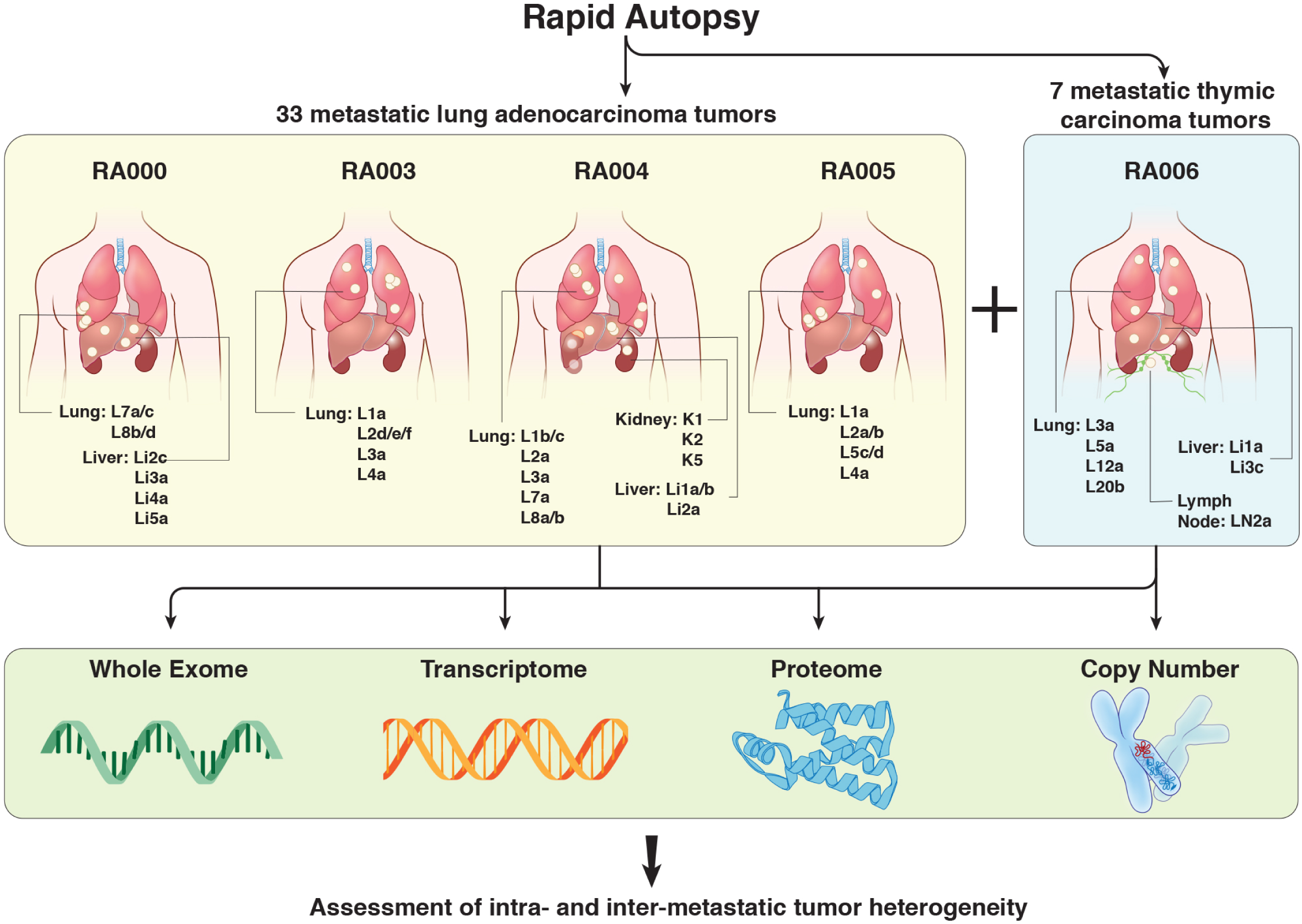
Flowchart of Study. Five patients underwent rapid autopsy (defined here as within 3 hours of patient death). Thirty-three metastatic lung adenocarcinoma (LUAD) and 7 metastatic thymic carcinoma tumors from lung, liver and kidney were subjected to whole-exome sequencing (WES), RNA-sequencing, copy number analysis, and mass-spectrometry based proteomics followed by assessment of intra- and inter-metastatic tumor heterogeneity. Three samples were removed from the study after sequencing due to low tumor content.

### Intra- and inter-metastatic mutational tumor heterogeneity is highly variable within patients and can be extreme

We performed whole-exome sequencing (WES) on the metastatic tumors, primary tumors, where available, and matched germline DNA from each patient. A range of 182 (RA005) to 1058 (RA003) non-silent mutations were identified per patient (Supplementary Table 2). RNA-seq demonstrated a high, independent validation rate of WES (Supplementary Fig. 2), similar to previous studies^19^. We found significant mutational heterogeneity within each patient, with the percentage of non-truncal variants ranging from 67% in RA000 to 99% in RA003 (Fig. 2a).

**Figure 2:**
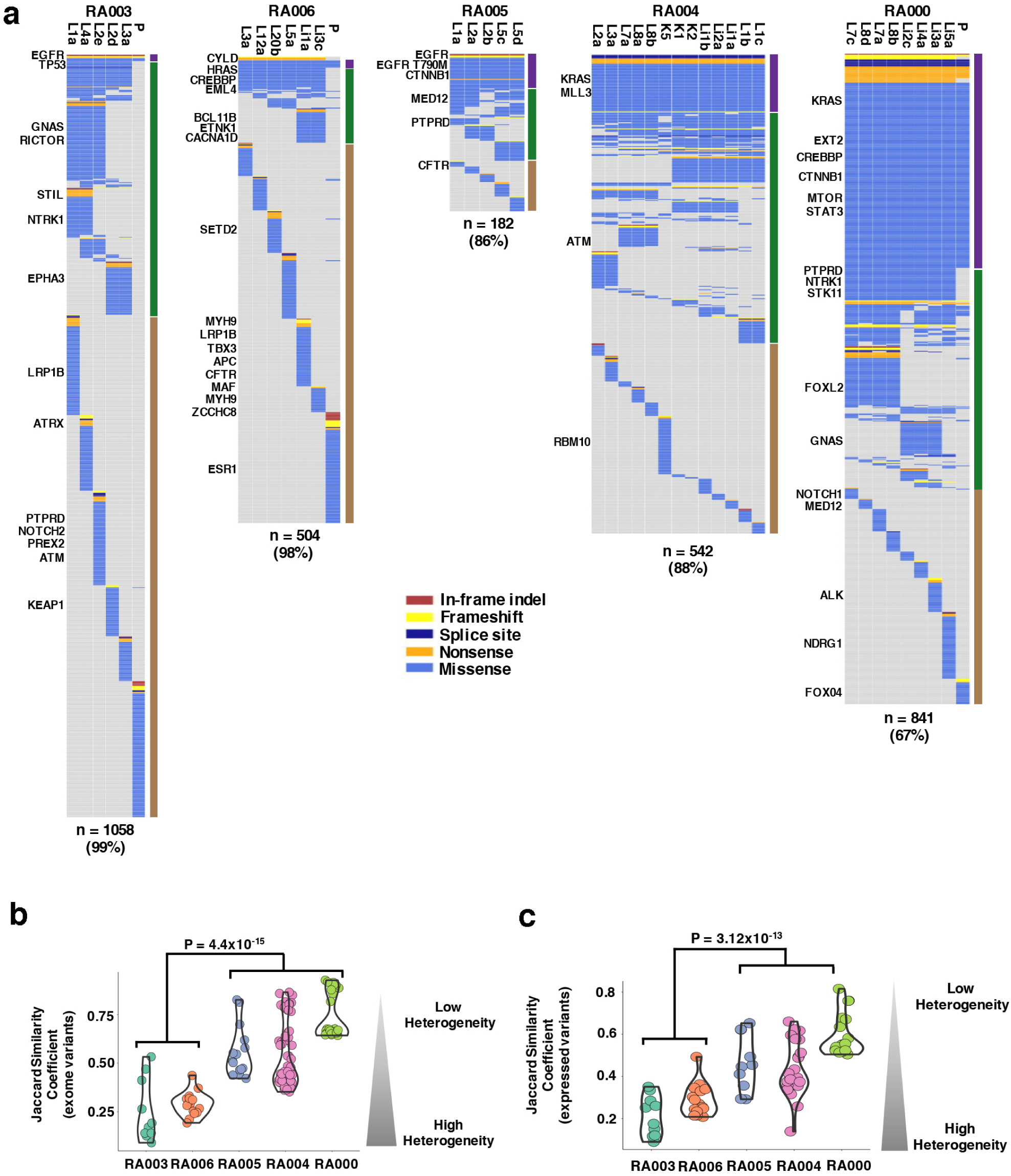
Intra- and inter-metastatic heterogeneity of somatic mutations of tumors from all autopsy patients. **(a)** Heat maps depict the distribution of non-silent somatic mutations among metastatic, and where available, primary tumors for each patient. Driver mutations are listed to the left of each heat map. The total number of non-silent mutations and the percentage of non-truncal mutations are shown below each heat map. The bars to the right of each heat map summarizes intra- and inter-metastatic heterogeneity; mutations present in all regions (purple), in more than one but not all (green), or only in one region (brown). Jaccard similarity coefficients of metastases within each patient based on mutations identified by **(b)** exome sequencing and **(c)** expressed variants by RNA-seq. Each circle represents the Jaccard similarity coefficient between two metastases. Coefficients range from zero to one representing highest and lowest heterogeneity, respectively. The P-value for the difference in mean Jaccard similarity coefficients between two groups of patients is shown. P: tumor sample obtained at diagnosis; L: lung tumor at autopsy; Li: liver tumor at autopsy; K: kidney tumor at autopsy

Activating mutations in *EGFR* (RA003), and *KRAS* (RA000, RA004) were present in all tumors – primary and metastatic. However, an activating *HRAS* mutation was present in all metastatic sites of patient RA006 at autopsy, but not in the primary tumor at the time of diagnosis. Patients RA003 and RA006 had an average of 12 and 14 non-truncal driver mutations, respectively. In contrast, there was an average of only 3.7 non-truncal driver mutations in the other three patients (RA005, RA004, and RA000), demonstrating variability in driver mutation acquisition among patients (Fig. 2a). Intra-tumor mutational heterogeneity in driver mutations was found when tested in RA003 tumors L2d/L2e (Fig. 2a).

We next calculated Jaccard similarity coefficients (defined as the ratio of shared to all mutations between two metastatic tumors) for each patient to quantitatively assess intra- and inter-metastatic tumor genomic heterogeneity (values 0 - 1 correspond to the range from minimal to maximal heterogeneity^17,20^) (Supplementary Table 3). The means of the Jaccard similarity coefficients for each patient ranged from 0.25 (RA003) to 0.73 (RA000) and were significantly different between patients (P=2.2×10^−16^, Kruskal-Wallis rank sum test, Fig. 2b). Two patients, RA003 and RA006, were clear outliers and exhibited what we termed “extreme” mutational heterogeneity. These two patients had significantly lower combined mean Jaccard similarity coefficients compared to the other patients (mean 0.28 vs. 0.57, P=2×10^−16^, Chi squared test) (Fig. 2b). Jaccard similarity coefficients exhibited a similar trend based on expressed mutations identified by RNA-seq analysis (Fig. 2c). Additional sequencing resulting in a median exome coverage of 487x (range 435 to 528) for the tumors from patients RA003 and RA006 did not substantially alter the observed extreme level of heterogeneity (Supplementary Fig. 3 and Supplementary Table 3). Collectively, these results indicate that intra- and inter-metastatic mutational heterogeneity can vary considerably among patients, with extreme heterogeneity evident in a subset of patients.

### APOBEC mutagenesis strongly correlates with mutational tumor heterogeneity

We next analyzed mutational signature profiles for each tumor and generated phylogenetic trees layered with mutational signatures^1,2^ to elucidate whether specific mutational processes could explain the observed variability in mutational tumor heterogeneity. Smoking signature mutations (C->A) were highly prevalent in patients RA000 and RA004, who were smokers (Supplementary Fig. 4, c). Mutations generated by the cytosine deaminase activity of APOBEC (apolipoprotein B mRNA editing enzyme, catalytic polypeptide-like) family of enzymes (substitutions of C with T or G in TCA or TCT motifs^21^) were most prevalent in patients RA003, RA005 and RA006, who were all non-smokers (Supplementary Fig. 4, d, e). Smoking signature mutations were generally truncal whereas APOBEC-signature mutations were largely in the shared and private branches of the phylogenetic trees (Fig. 3a-e).

**Figure 3:**
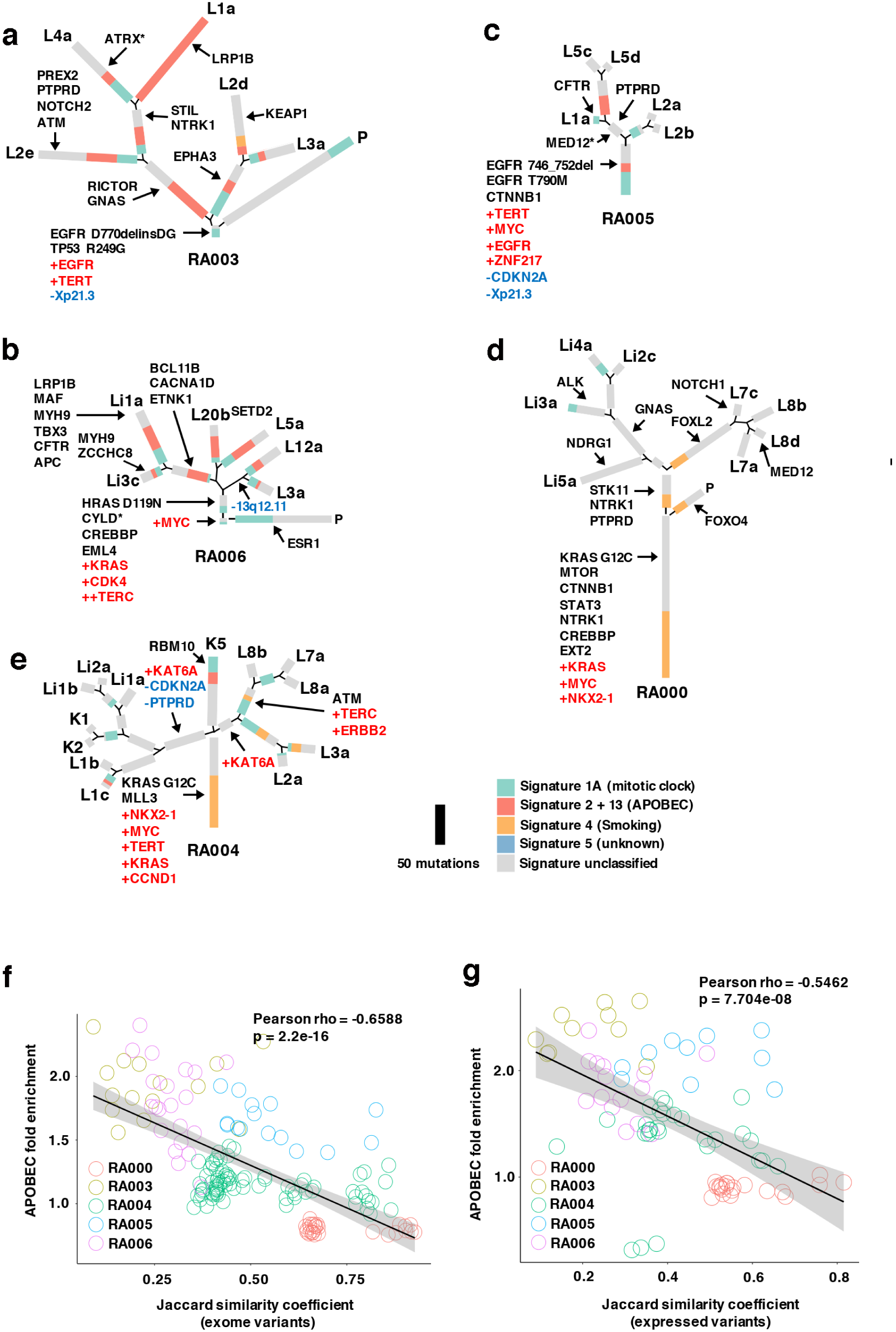
Inferred phylogeny, mutational signatures and APOBEC associated heterogeneity. Phylogenetic trees were generated from all validated mutations identified by whole-exome sequencing from tumors within patient **(a)** RA003, **(b)** RA006**, (c)** RA005, **(d)** RA000, and **(e)** RA004 using the maximum parsimony method. Trees are rooted in mutations common to all tumors within each patient. Trunk and branch lengths are proportional to the numbers of mutations acquired on the corresponding trunk or branch. Each private branch represents mutations unique to each individual tumor. Colors represent COSMIC mutational signatures. Driver mutations and focal copy number amplifications/deletions are mapped to the trunks and branches as indicated. Asterisks denote nonsense mutations. APOBEC fold enrichment correlates with **(f)** DNA Jaccard similarity coefficients and **(g)** RNA Jaccard similarity coefficients across tumors of all autopsy patients. Each circle represents mean APOBEC fold enrichment and the Jaccard similarity coefficient between two tumors from a given patient. P: tumor sample obtained at diagnosis; L: lung tumor at autopsy; Li: liver tumor at autopsy; K: kidney tumor at autopsy.

To further assess the timing of APOBEC-induced mutagenesis, we evaluated mutational signatures in available tumors collected at the time of diagnosis. We found no evidence of APOBEC mutagenesis in these samples, including those from patients RA003 and RA006 (Fig. 3a, b), indicating that APOBEC-signature mutations were acquired later, either during further progression of metastatic disease or upon subsequent treatment. We next evaluated the relationship between APOBEC mutagenesis and heterogeneity. APOBEC mutation fold enrichment, a measure of APOBEC mutagenesis^21^ (Supplementary Table 4), strongly correlated with Jaccard similarity coefficients based on WES variants (Pearson rho = −0.66, p = 2.2e-16) and expressed variants by RNA-seq (Pearson rho = −0.55, p = 7.704e-08) (Fig. 3f, g). Taken together, these results suggest APOBEC mutagenesis significantly contributes to the generation of intra- and intermetastatic mutational heterogeneity.

### Expression of *APOBEC3* region transcripts correlates with APOBEC mutagenesis

Next, we examined RNA-seq data to determine whether *APOBEC3* region transcript expression contributes to variability in APOBEC mutagenesis. Among the LUAD patients, we found *APOBEC3B* to be expressed at higher levels than *APOBEC3A*; this was particularly evident in the tumors of patient RA003 (Fig. 4a). APOBEC mutagenesis (measured here by the counts of t<u>C</u>w to t<u>T</u>w and t<u>G</u>w mutations)^21^ was highly correlated with *APOBEC3B* expression (Pearson rho=0.68, P=9.4×10^−05^) but not with *APOBEC3A* expression (Pearson rho =0.19, P=0.34) (Supplementary Fig. 5, b). Analysis of isoform-specific expression of *APOBEC3A* and *APOBEC3B* with custom TaqMan assays confirmed these results (Supplementary Fig. 5, d). Moreover, within patient RA003, who displayed multiple tumors with significant APOBEC mutagenesis (Fig. 4a), *APOBEC3B* was expressed 20 to 50-fold higher than *APOBEC3A* (Fig. 4b).

**Figure 4:**
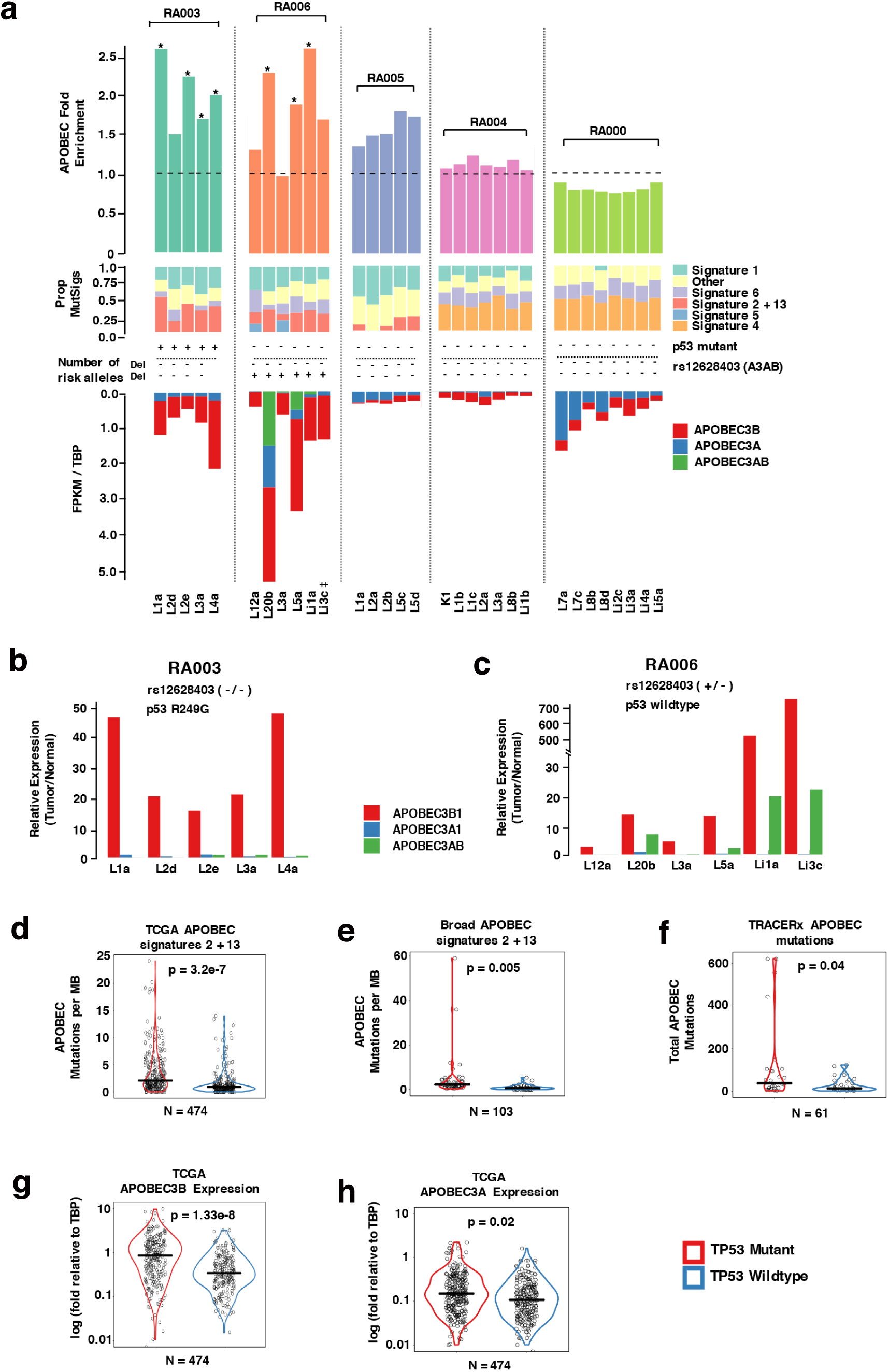
Relationship between APOBEC fold enrichment, mutational signatures, *APOBEC3* region transcript expression, *APOBEC3AB* germline variant and mutant *TP53*. **(a)** APOBEC fold enrichment of each tumor. Asterisks denote significant enrichment. Dashed line denotes zero enrichment of APOBEC mutations. Proportion of COSMIC signatures for each tumor appear below APOBEC fold enrichment. Number of risk alleles for *APOBEC3* germline variant rs12628403 is shown below the proportion of COSMIC signatures. Expression of *APOBEC3B, APOBEC3A* and *APOBEC3AB* are shown below the proportion of COSMIC signatures. Relative expression of isoforms *APOBEC3B1, APOBEC3A1* and *APOBEC3AB* in tumor relative to normal lung or liver by custom TaqMan assays in **(b)** patient RA003 and **(c)** patient RA006. Six of the 37 tumors are not included here due to insufficient RNA for sequencing. **(d)** APOBEC signature mutations per megabase in TCGA and **(e)** Broad dataset and **(f)** total APOBEC mutations in TRACERx dataset in *TP53* mutant compared to *TP53* wildtype tumors. **(g)** *APOBEC3B* expression and **(h)** *APOBEC3A* expression in *TP53* wildtype and mutant tumors in the TCGA dataset. P-values shown are adjusted for patient age and number of pack years smoked. ‡-visual inspection of RNA-seq data shows expression of *APOBEC3AB*; see Supplementary Note for details.

To elucidate factors affecting APOBEC mutagenesis in our patients, we genotyped an *APOBEC3* germline variant, rs12628403, associated with increased APOBEC mutagenesis^22^. This germline variant is a proxy for a 30-kb deletion that fuses the coding region of *APOBEC3A* with the 3’ UTR of *APOBEC3B* to generate a chimeric *APOBEC3A-APOBEC3B (APOBEC3AB)* transcript. This chimeric *APOBEC3AB* transcript is more stable than the *APOBEC3A* transcript and leads to higher APOBEC3A protein levels *in vitro^23^*. Only patient RA006, the thymic carcinoma patient, was a carrier of the rs12628403 allele, and was predicted to generate the *APOBEC3AB* transcript (Fig. 4a). Indeed, we found expression of *APOBEC3AB* transcripts in patient RA006 tumors from RNA-seq data (Fig. 4a) and validated expression of this transcript using a TaqMan assay (Fig. 4c). However, the *APOBEC3B* transcript was also expressed in tumors from patient RA006, suggesting that both may have contributed to APOBEC mutagenesis. Expression of *APOBEC3B* (Pearson rho=0.77, P=0.08) and *APOBEC3AB* (Pearson rho=0.62, P=0.18) but not *APOBEC3A* (Pearson rho=0.23, P=0.67) significantly correlated with APOBEC mutagenesis (Supplementary Fig. 6-c).

Although expression of *APOBEC3AB* was lower than *APOBEC3B* (Fig. 4c), APOBEC3A encoded by *APOBEC3AB* is considered a more potent inducer of mutagenesis than APOBEC3B^23,24^. Therefore, we quantified the contribution of APOBEC3A and APOBEC3B to APOBEC mutagenesis by calculating YTCA and RTCA enrichment (where Y is a purine and R is a pyrimidine), attributed to differential activity of these enzymes^25^. In all tumors with high APOBEC mutagenesis from patient RA006, there was significant enrichment of YT<u>C</u>A compared to RT<u>C</u>A (Supplementary Fig. 6), thereby suggesting APOBEC3A-like mutagenesis as a likely driver of extreme heterogeneity in this metastatic thymic carcinoma patient with a germline *APOBEC3AB* deletion. Together, our results implicate expression of *APOBEC3* region transcripts as a mediator of APOBEC mutagenesis in metastatic lung adenocarcinoma and thymic carcinoma.

#### *TP53* mutations are associated with APOBEC hypermutators in lung adenocarcinoma

We hypothesized that mutant *TP53*, present only in LUAD patient RA003, may be contributing to high *APOBEC3B* expression and increased APOBEC mutagenesis. Using three independent LUAD datasets^19,26,27^, we found *TP53* mutations correlated with higher counts of APOBEC-signature mutations (Fig. 4d-f). Moreover, mutant *TP53* was associated with APOBEC ‘hypermutators’^22^ in our cohort and in all three datasets (Table 1). Mutant *TP53* was also associated with significantly higher expression of *APOBEC3B*, as well as an increase in *APOBEC3A*, compared to wild-type *TP53* tumors in the TCGA dataset (Fig. 4g, h). Thus, our results suggest mutant *TP53* contributes to increased *APOBEC3B* expression, APOBEC mutagenesis and is associated with APOBEC hypermutators in lung adenocarcinoma.

**Table 1:**
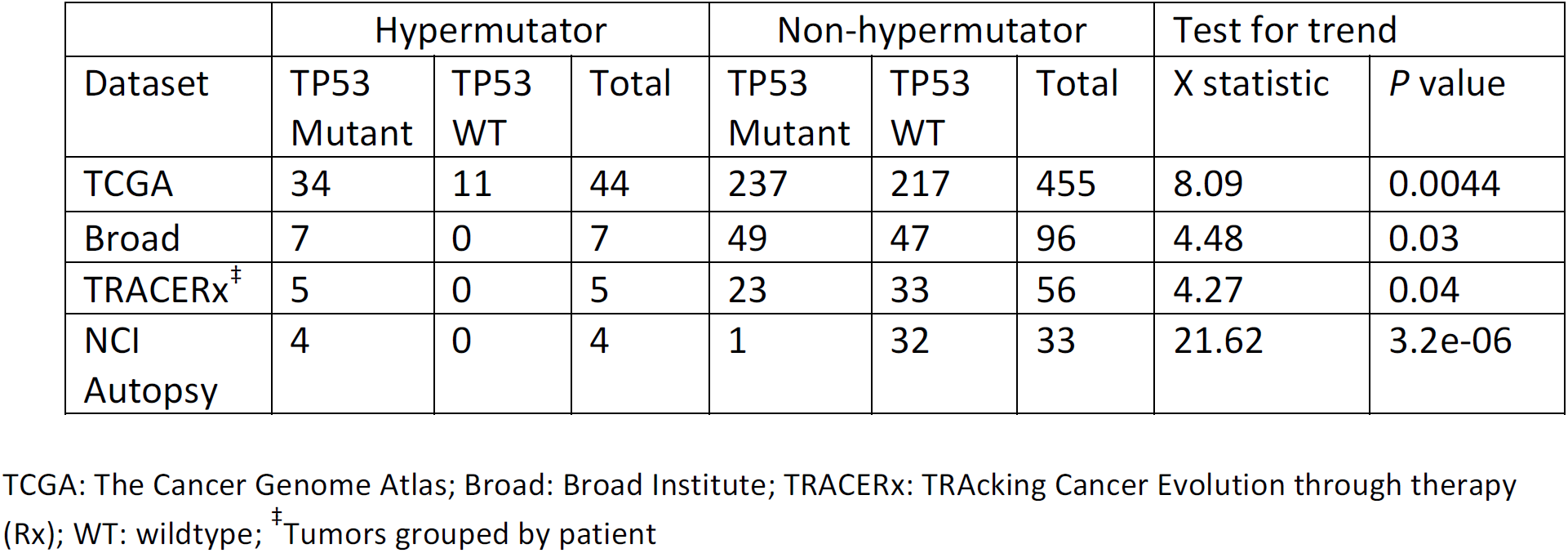
Relationship between TP53 mutation status and APOBEC hypermutators in lung adenocarcionoma

### Integration of copy number, transcript and protein abundance highlights mechanisms of proteomic heterogeneity

To evaluate multi-dimensional heterogeneity, we plotted Pearson correlation coefficients (PCCs) for each data type between pairs of tumors for each patient across all genes for which copy number, transcript expression, and protein abundance data were available (Fig. 5a-e). Each patient displayed variable patterns of heterogeneity across each data type. Patient RA003 exhibited the least (Fig. 5a), whereas patient RA004 the most (Fig. 5d) heterogeneity. Patient RA005 showed the most heterogeneity in gene expression and protein abundance only between tumor L5d and tumors L2a/L2b/L5c (Fig. 5c). Patient RA006 showed lower heterogeneity in copy number, gene expression and protein abundance within three pairs of tumors (L20b/L5a, Li3c/Li1a, L12a/L3a) compared to other pairs (Fig. 5b).

**Figure 5:**
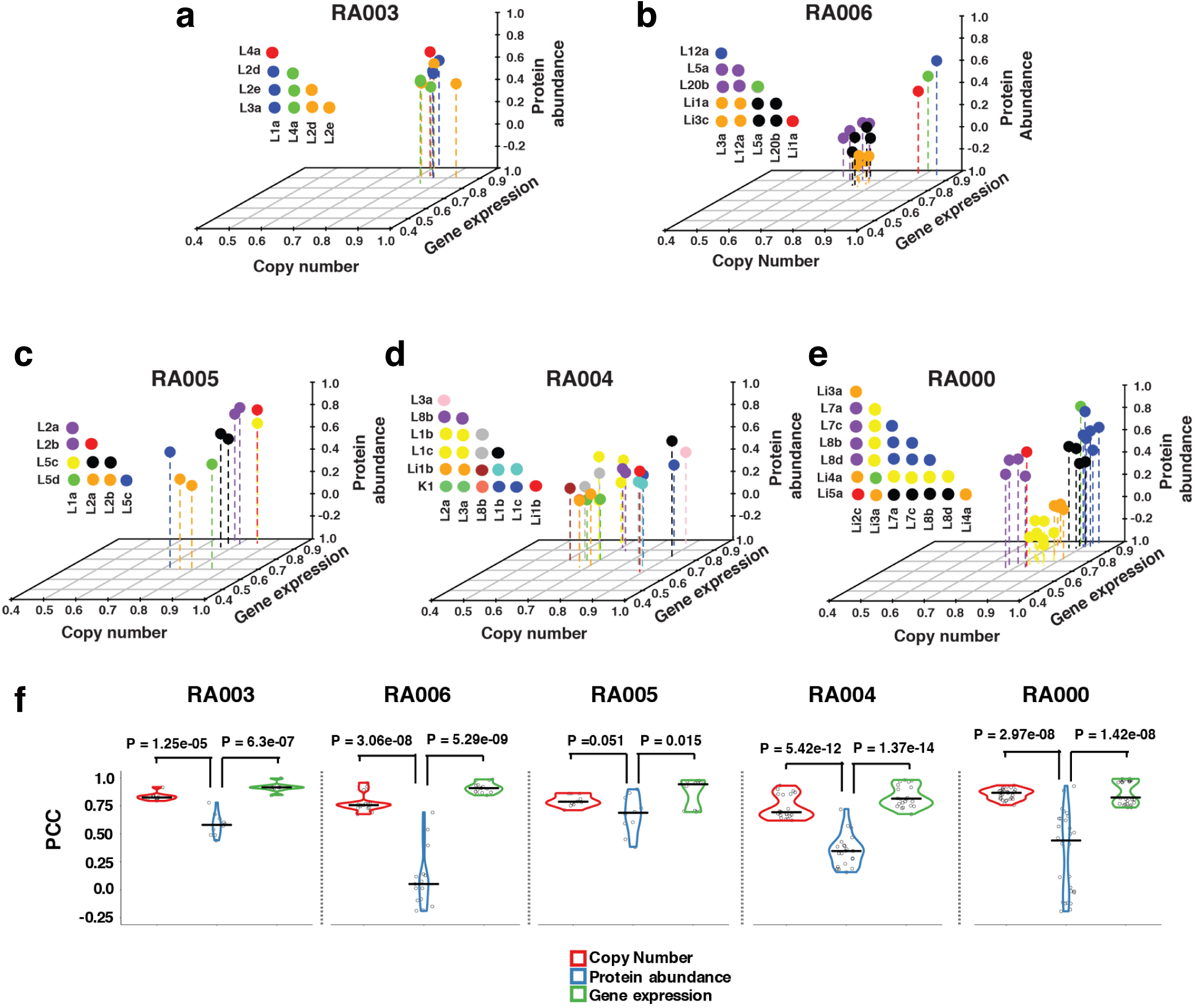
Intra-patient multi-omic tumor relationships. Pearson correlation coefficients (PCCs) are shown on each axis across all common genes identified in copy number, gene expression and protein abundance datasets between all tumors within patient **(a)** RA003, **(b)** RA006, **(c)** RA005, **(d)** RA004, and **(e)** RA000. Each circle represents the relationship between two tumors. Previously designated tumor clusters are assigned the same color. Colors are assigned independently for each patient. **(f)** PCCs between tumors from each patient grouped by data type: copy number, gene expression and protein abundance. P-values for the difference in mean PCCs between data types are shown for each patient.

Next, we plotted PCCs within each patient for each data type. Protein heterogeneity was significantly greater than gene expression and CNA heterogeneity within all patients (Fig. 5f). We then performed pairwise comparison of the PCCs of each data type (Supplementary Fig. 7). We found a strong, positive linear correlation between CNA heterogeneity and protein heterogeneity for patients RA004 (Pearson rho=0.68, p=7.5e-04) and RA006 (Pearson rho=0.80, p=3.74e-4) but not for the other patients (Supplementary Fig. 7, d, g, j, m), providing evidence that CNAs can lead to protein heterogeneity in these patients. Gene expression heterogeneity was also associated with protein heterogeneity but only in patients RA005, RA006 and RA000 (Supplementary Fig. 7, k, n). Together, these results demonstrate high heterogeneity in protein abundance between metastases of these patients which could stem from heterogeneity in CNA and gene expression.

### Late-event CNAs contribute to heterogeneity in gene expression and protein abundance

We next performed hierarchical clustering by chromosomal cytoband, gene expression and protein abundance to further evaluate heterogeneity within each data type. Metastases from each patient clustered together for CNA, gene expression and protein abundance (Fig. 6a, Supplementary Fig. 8, 9a-b). Metastases from patients RA004 and RA006 showed the lowest correlation in protein abundance (Supplementary Fig. 9, d) and clear differences in CNAs (Fig. 6a and Supplementary Table 5). To explore the downstream effects of CNAs, we plotted gene expression and protein abundance ratios of genes within each chromosomal arm between metastatic lineages of each patient (Fig. 6b, c and Supplementary Tables 6 and 7). Arm-level CNAs within tumors of patients RA006 and RA004 corresponded with changes in gene expression and protein abundance of genes at the corresponding arm-level. For example, copy number differences in arm 4p between RA006 tumors corresponded with changes in expression and protein abundance (Fig. 6b). Where there were no copy number differences, such as in arm 7q in patient RA006 and arms 4p/7q of patient RA003, there were no corresponding changes in gene expression and protein abundance (Fig. 6c).

**Figure 6:**
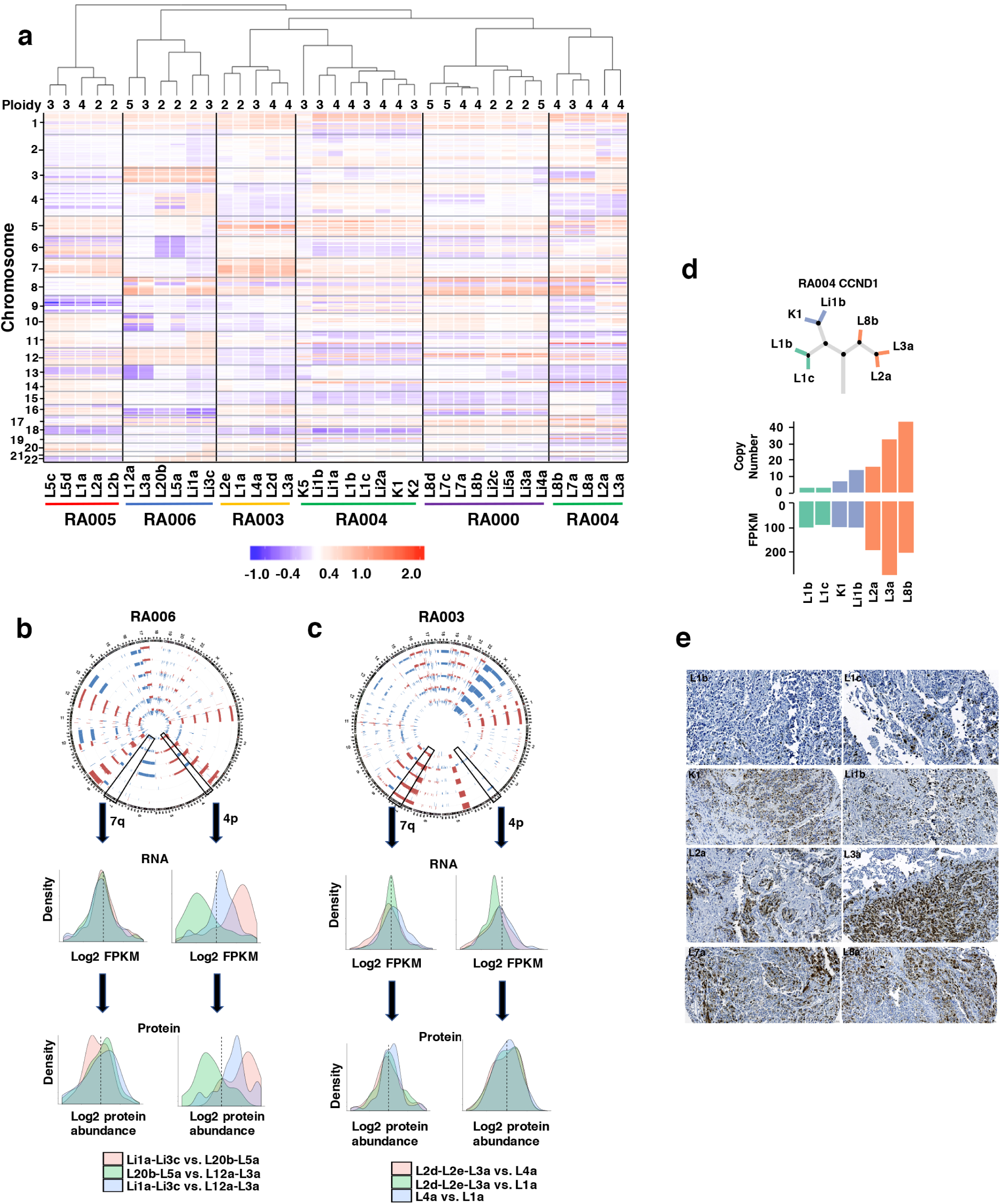
Copy number heterogeneity corresponds with transcriptomic and proteomic heterogeneity. (**a**) Hierarchical clustering by copy number across the genome across all tumors from all patients (at cytoband resolution). Losses (purple) and gains (red) in log2 scale are depicted relative to mean ploidy. Mean ploidy is shown in the top row (rounded to nearest integer). Copy number differences in chromosomal arms 4p and 7q between tumors of (**b)** patient RA006 and **(c)** patient RA003 and corresponding changes in FPKM and protein abundance. Probability density plots show log ratios of mean FPKM by chromosomal arm for sets of tumors as displayed. X-axis dashed line denotes ratio of 1 (or log ratio 0). For visualization purposes, X-axis was cut at log −1 and 1 for RNA and log −0.1 and 0.1 for Protein. Y-axis was cut at probability density of 2. **(d)** Phylogenetic tree depicts tumors of patient RA004 with corresponding copy number and RNA-seq FPKM of CCND1 for each tumor. **(e)** Protein expression of CCND1 for tumors of patient RA004 as assessed by immunohistochemistry from tissue microarrays.

Heterogeneity in focal-level CNAs also corresponded with changes in RNA and protein. For example, *CCND1* was highly amplified in patient RA004 tumors L2a, L3a and L8b (Supplementary Table 8) which corresponded with high gene and protein expression. On the other hand, tumors L1b and L1c in which there was minimal increase in *CCND1* copy number, gene and protein expression were low (Fig. 6d, e). Interestingly, among patient RA004 liver and kidney tumors, Li1b and K1, *CCND1* was highly amplified with correspondingly high protein but moderate gene expression, suggesting tissue specific discordance of gene and protein expression of select genes (Fig. 6d, e).

We next constructed phylogenetic trees based on CNAs for each patient. Both arm and focal-level CNAs largely occurred early in tumor development (i.e. truncal) in patients RA000, RA003 and RA005 (Supplementary Fig. 10, 11, 12 a-b) but occurred later in tumor development (i.e. shared and private) in patients RA004 and RA006 (Fig. 3, Supplementary Fig. 13 and 14 a, b). These results suggest that late, not early, CNAs likely contributed to the observed changes in gene expression and protein abundance between metastases of these patients. Differential focal and arm-level CNAs may reflect ongoing chromosomal instability as well as selective pressure during evolution of metastatic lineages.

### Enrichment of interferon signaling pathways in tumors with high *APOBEC3* expression and immune heterogeneity

We next sought to decipher any common gene sets or pathways within the RNA-seq and mass-spectrometry proteomics data that were heterogeneously enriched within each patient. Using an unbiased approach with single-sample gene set enrichment analysis (ssGSEA)^28,29^, we found interferon (IFN)-signaling pathways (related to activity of IFN-α, IFN-β and IFN-γ), were the most significantly and differentially enriched pathways within patient RA003 (tumors L1 and L4a) and RA006 (tumors L20b, L5a, Li1a and Li3c) (Fig. 7a-d) at both gene expression and protein abundance levels. No common outlier gene sets were identified for the negatively enriched pathways. Within the tumors from patients RA000, RA004 and RA005, no recurrent, common pathways were identified by GSEA of both the transcriptome and proteome (Supplementary Table 9). The six tumors from patients RA003 and RA006 that were enriched in IFN-signaling pathways also had the highest expression of *APOBEC3* region transcripts (Fig. 3a, b, c). Given that *APOBEC3A* and *APOBEC3B* are IFN-stimulated genes^30–32^, our results suggest IFN-signaling within the tumor immune microenvironment as a potential mechanism of heterogeneity in *APOBEC3* region transcript expression.

**Figure 7:**
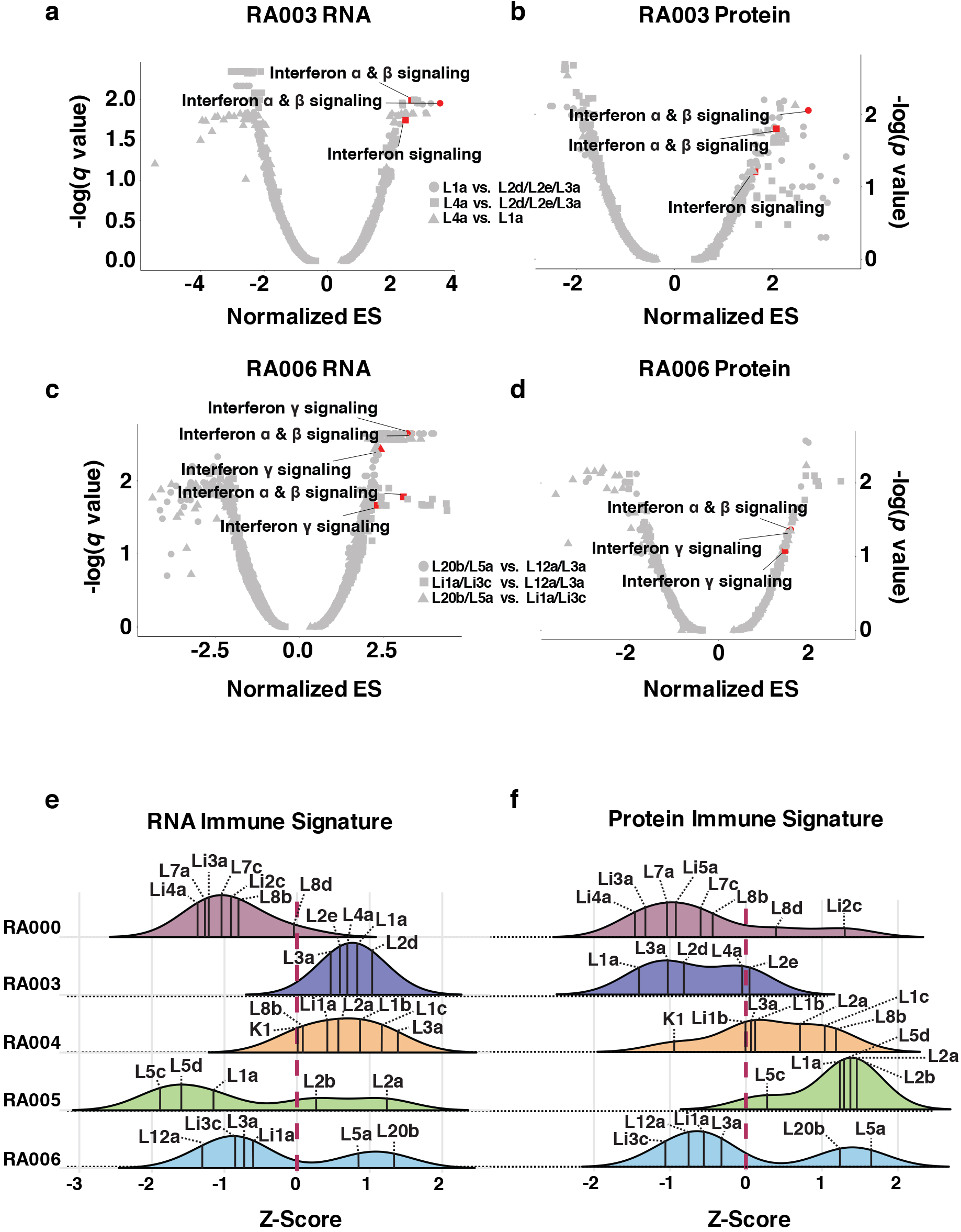
Site specific enrichment of interferon signaling pathways and immune signature heterogeneity. Single-sample gene set enrichment (ssGSEA) analysis of transcriptome and proteome using the REACTOME databases are shown for tumors from patients RA003 **(a, b)** and RA006 **(c, d)**. Significantly enriched interferon pathways (q < 0.05 for transcriptome, p < 0.10 for proteome) are colored red. Immune signature scores within the **(e)** transcriptome and **(f)** proteome are shown between all tumors of patients RA000, RA003, RA004, RA005 and RA006. Scores were normalized across all tumors separately for transcriptome and proteome. ES: enrichment score. Symbols denote comparisons between groups of tumors as indicated.

To further interrogate heterogeneity in the immune microenvironment between tumors of each patient in an unbiased manner, we analyzed the gene expression and protein abundance data using ssGSEA based on CIBERSORT immune genes^33^. The overall immune signature score for tumors within each patient varied considerably between transcriptome and proteome in patients RA003 and RA005 (Fig. 7e, f). Patients RA000 and RA004 showed low and high overall immune signature scores, respectively (Fig. 7e, f). In contrast, tumors from patient RA006 showed large and consistent differences in immune signature scores across both the transcriptome and proteome (high in L20b and L5a vs. low in L12a, L3a, Li1a, Li3c), demonstrating heterogeneous immune cell infiltration in the tumor microenvironment of this patient.

## DISCUSSION

Genomic, transcriptomic and proteomic analyses of tumors from multiple anatomic sites sampled through rapid autopsy offer a unique opportunity to comprehensively explore the biological processes that shape the evolution of metastatic tumors. Here, through rapid autopsy, we have characterized, for the first time, the proteogenomic evolution of metastatic lung and thymic carcinoma through exome and transcriptome sequencing, CNA analysis, and unbiased quantitative mass spectrometry-based proteomics of 37 metastatic tumors. Most importantly, we have uncovered mechanisms likely driving the mutational, transcriptomic and proteomic landscape of these metastatic tumors.

At the genomic level, we provide evidence that APOBEC mutagenesis may be a driver of mutational heterogeneity in metastatic lung and thymic carcinoma tumors. APOBEC mutagenesis has been described as one of the most common mutational processes second only to “ageing”^34^. To date, however, within thoracic tumors, this process has been described mostly in primary tumors^1,35,36^. These studies have shown APOBEC mutagenesis to be associated with subclonal mutations that occur late in the evolution of primary tumors and within spatially distinct regions^1,2^. In our study, we identified a subset of patients with high APOBEC-mutagenesis in metastases, but not in the primary tumor, suggesting APOBEC-mutagenesis can generate mutations late in the evolution of metastatic disease. Given that all patients in our cohort received prior treatment, we cannot exclude the possibility that therapy contributed to the observed findings. However, recent studies assessing tumors pre- and post-chemotherapy in multiple tumor types did not find an increase in overall mutational load or a significant increase in APOBEC-signature mutations^37,38^. Moreover, the patients in our cohort with the highest level of APOBEC-mutagenesis, RA003 and RA006, received the least therapy prior to autopsy.

While the existence of mutational heterogeneity in metastases has been previously described^10,12^, the mechanisms have not been clear^39^. Our data suggest APOBEC mutagenesis can generate both putative driver and passenger mutations late in metastases, thereby generating inter-metastatic mutational heterogeneity that in some cases can be extreme. These results stand in contrast to recent genomic studies of the metastases of patients with pancreatic^17^ and prostate^16^ cancer, which have shown limited mutational heterogeneity and no significant APOBEC mutagenesis^21^. Both of these tumor types have also shown no evidence of APOBEC mutagenesis within primary tumors highlighting the likely histologic specificity of this process. Ultimately, the clinical importance of APOBEC mutagenesis will be determined by the response of heterogeneous metastatic tumors - with and without APOBEC-signature mutations - to chemotherapy, targeted agents and/or immunotherapy.

Both APOBEC3A and APOBEC3B have been shown to localize to the nucleus^40^ leading to potent DNA damage^41^, deaminase activity and base substitutions in the genome^35,42^. Upregulation of *APOBEC3B* causes APOBEC-signature mutations *in vitro*^43^. Expression of *APOBEC3A* and *APOBEC3B* has also been associated with APOBEC-signature mutations in primary tumors from multiple cancer types including LUAD^25,35,36,44^. However, it is unclear whether these same transcripts promote APOBEC mutagenesis in metastatic lung and thymic carcinoma tumors. In our set of metastatic tumors, we observed a strong correlation between expression of *APOBEC3B* and *APOBEC3AB* transcripts and APOBEC mutagenesis suggesting expression of such transcripts may be a more dominant mechanism of APOBEC mutagenesis in metastatic thoracic tumors as opposed to earlier stage disease. These results are in line with recent work in breast cancer that has shown higher APOBEC mutagenesis in metastatic disease compared to early stage primary tumors^45^.

Mutant *TP53* has previously been associated with higher APOBEC-signature mutations in breast cancer^35^ and there is recent evidence that TP53 can repress *APOBEC3B* expression through direct transcriptional regulation of its promoter^35,46,47^. Our results suggest mutant *TP53* may be an important contributor of increased *APOBEC3B* and subsequent generation of APOBEC-signature mutations in LUAD. In particular, we show mutant *TP53* is associated with APOBEC hypermutators in LUAD using three large-scale independent datasets. Whether such APOBEC hypermutators also display extreme mutational heterogeneity similar to patient RA003, and how these patients may respond to therapy will be important to assess in future clinical trials.

Expression of the *APOBEC3AB* transcript was captured in our study by the presence of the *APOBEC3* germline variant, rs1262840, within thymic carcinoma patient RA006. This germline variant has previously been associated with increased APOBEC-signature mutations in primary breast cancer tumors^22^. Whether this variant is also associated with high APOBEC-mutagenesis and mutational heterogeneity within thymic carcinoma is unknown, as no previous multi-region sequencing study has been performed on this rare tumor type. Additionally, to our knowledge, apart from the current study, this variant has not been examined in relation to APOBEC-mutagenesis in metastatic disease. Given that this germline variant can easily be tested utilizing blood DNA, our results warrant further testing of the association between this *APOBEC3* germline variant with APOBEC mutagenesis and mutational tumor heterogeneity in thymic carcinoma as well as other metastatic tumor types.

Our integrated CNA, RNA-seq and quantitative mass spectrometry study demonstrate that late-event CNAs can be important drivers in the evolution of metastatic cancer through downstream changes in transcript and protein abundance. Early studies in yeast showed CNAs for a given gene lead to proportional increases in protein abundance^48–51^. More recently, studies in primary tumors demonstrated variability in CNA to protein cis-effects^52–54^. In metastatic disease, multiple studies have reported late-event CNAs^12,18,55–58^ but the effect of CNAs on transcript and protein abundance has not been examined.

In one recent metastatic pancreatic cancer study, CNA differences among tumor suppressor genes were not evident at the protein level by IHC suggesting late-event CNAs can be stochastic changes rather than evolutionary selected events^17^. In the current study, all patients had some evidence of late-event CNAs. However, only patients with significant differences in late-event CNAs between tumors exhibited corresponding differences in transcript and protein abundance. In light of the recent association between copy number heterogeneity and increased recurrence and death in early stage NSCLC^2^, our results raise the question of whether late-event CNAs, through downstream effects on gene expression and protein abundance, can result in worse outcomes for a subset of patients with metastatic cancer. Proteomic heterogeneity induced in part by CNAs may also explain why chromosomal instability (CIN) has been associated with poor outcomes in cancer^2,59–61^. Importantly, proteomic heterogeneity in our set of metastatic tumors, was much higher than CNA and transcriptomic heterogeneity, suggesting other mechanisms such as epigenetic and post-translational modifications may also be important drivers of proteomic heterogeneity.

Through our unbiased analysis of transcriptomic and proteomic data, we found enrichment of IFN signaling within the microenvironment of tumors of patients with the highest *APOBEC3* region transcript expression. *APOBEC3A* and *APOBEC3B* are IFN-stimulated genes induced *in vitro* by IFN stimulation and viral infections that activate an IFN response^30–32^. These data support the known role of APOBEC mutagenesis contributing to non-cytolytic viral clearance^62^. Our data suggests IFN signaling within the tumor microenvironment may, in part, influence *APOBEC3* region transcript expression and thereby contribute to heterogeneity in APOBEC-signature mutations within the tumors of a given patient.

We also found transcriptomic and proteomic heterogeneity in immune signatures within and between patients. Major advances have been made in the treatment of metastatic tumors, including lung adenocarcinoma and thymic carcinoma, through immunotherapies such as immune checkpoint blockade^63,64^. Nonetheless, only a subset of patients responds and metastases within a given patient may respond differently due to immune heterogeneity^65^. Even without immunotherapy, metastases within a patient may also have differing tumor immune microenvironments, as we demonstrate within thymic carcinoma patient RA006 and as has been recently shown within an ovarian cancer patient^66^. We further demonstrate that, within a given patient, the tumor immune microenvironment may exhibit substantial differences between the transcript and protein expression, adding to the complexity of assessing the immune microenvironment.

One strength of our study is the comprehensive examination of tumor heterogeneity by integrating genomic (exome, CNA), transcriptomic (RNA-seq) and proteomic (global mass spectrometry analysis for protein abundance) data from multiple metastatic tumors procured through rapid autopsy. One of the limitations of our study is that all autopsy patients in this study were diagnosed at the late stage of metastatic disease. Hence, we were unable to conduct a complete temporal analysis of tumor evolution from early to late stage disease. The ongoing TRACER_x_ study^2,26^ and co-recruitment of those patients to the PEACE (Posthumous Evaluation of Advanced Cancer Environment) post-mortem study^67^ will allow for better elucidation of the evolution primary tumor to metastatic advanced disease, including at the end of life.

In conclusion, in this report, we present the heterogeneous genomic, transcriptomic and proteomic landscape of metastatic lung and thymic carcinoma as well as identify possible mechanisms underlying such multi-level heterogeneity. High activity of the APOBEC3 enzymes, represented by transcript expression, and modulated by germline variants, mutant *TP53*, and the immune microenvironment, can greatly alter the genomic landscape between metastatic tumors of a given patient. Arm-level and focal CNAs occurring later in tumor evolution can generate significant downstream heterogeneity through effects on gene expression and protein abundance. Further studies by comprehensive analyses of multiple metastatic sites from larger patient populations, including different tumor types, are warranted to validate these mechanisms. Such an endeavor requires development of rapid autopsy programs, meticulous collection and processing of tumors from all possible sites of disease and integrated “omics” analyses. These tumor heterogeneity studies will be integral for evaluating the outcomes of ongoing clinical trials, developing new paradigms in clinical trial design and ultimately to improve survival for patients with metastatic cancer.

## METHODS

### Rapid Autopsy

Samples were obtained from five patients diagnosed with thoracic malignancies who underwent rapid autopsy. Informed consent for rapid autopsy was obtained under an IRB approved protocol 13-C-0131 (NCT01851395) entitled “A Pilot Study of Inpatient Hospice with Procurement of Tissue on Expiration in Thoracic Malignancies.” Patients previously treated at the NCI and with life expectancy less than 3 months were offered inpatient hospice treatment at the Clinical Center of the National Institutes of Health and upon death autopsies were initiated within 3 hours. One patient, RA003, elected to receive end of life care at home and was subsequently transported to the NIH Clinical Center post-mortem. Prioritization of lesions removed at autopsy was based on CT scan performed within one month before death. All tumors within each patient were removed by an experienced pathologist and macro dissected to remove surrounding non-neoplastic tissue. Punch biopsy needles were used to obtain spatially distinct cores from each tumor. One-third of each tissue core sample was fixed in 10% buffered formalin, one-third in optimal cutting temperature compound (OCT) and the remaining tissue was immediately flash frozen in liquid nitrogen and stored at −80 °C. For each tissue sample, a 5-μm section was taken to create a hematoxylin and eosin slide to visualize neoplastic cellularity using a microscope.

A normal control for each patient was represented by a normal tissue, if available, and/or a blood sample. DNA and RNA was isolated from approximately 30 mg of snap-frozen tumor tissue using the All Prep DNA/RNA Mini Kit (Qiagen). RNA was partially degraded with an average RNA integrity number 5.17 but was comparable between organs of different patients and similar in quality to previous post-mortem studies (van der Linden et al., 2014). To ensure adequate quality, samples RA003_L2f, RA006_LN2a, RA005_L4a were removed post-sequencing but before analyses due to low tumor content (less than 20 percent) based on Sequenza purity estimates.

### Whole-exome sequencing data processing, variants calling, filtering and annotation

Whole-exome sequencing of tumor and normal samples was performed at a sequencing core at the NCI Frederick National Laboratory at the National Cancer Institute (NCI). Libraries were constructed and then sequenced as 2 × 126 nt paired-end reads with Illumina HiSeq2500 sequencers. Mean coverage depth was 161x (range 114x to 231x). Raw sequencing data in FASTQ format were aligned against the reference human genome (hg19) with BWA^68^. The alignment BAM files were further processed following GATK’s best practices^69^ with Picard tools, namely MarkDuplicates, IndelRealigner, and BaseQRecalibrator. Somatic variants were then called from the processed BAM files using Strelka (v1.0.10)^70^ with the default version of BWA configuration file. The identified somatic variants reported in the “passed” vcf files by Strelka were used for further analysis. Variants were functionally annotated using snpEff/snpSift version 3.4 (see URLs) with databases of GRCh37.70 and dbNSFP version 3.4, and the types of variants were filtered using snpSift^71^. If a variant was also reported in one of the three public databases: 1000 Genome Project, ExAC, and ESP-NHLBI with a MAF greater than 5%, the variant was removed. For each patient, variants identified by Strelka^70^ from all tumor regions were combined to get a unique variant list. Using this patient-specific list, if a variant in a particular tumor site was not called by Strelka, the Samtools mpileup was used to retrieve the reference and alternative reads coverage for each SNV site. If the site had >= 2 alternative reads and VAF >=1%, the SNV was considered present in the tumor site. For short indels, if the variant site was not in the “passed” vcf file, the Strelka called “all” vcf file is used to retrieve the reference and alternative reads coverage; if missing in the “all” vcf file, the indel site was considered absent. Patient RA000 was previously known to have a KRAS G12C mutation based on molecular profiling. Although this mutation did not pass variant filtering, it was noted on manual review. Identified missense mutations were manually reviewed using the Integrative Genomics Viewer version 2.4^72,73^.

### Phylogenetic analysis

Phylogenetic analysis was conducted using Phangorn^74^ and phytools R packages with all identified variants (silent and non-silent) from all tumor sites in each patient after converting the mutation profile into binary format. The initial phylogenetic relationships between tumor regions for an individual patient was inferred using both the Maximum Parsimony and the Unweighted Pair Group Methods (UPGMA). Phylogenetic trees were then redrawn by hand in Adobe Illustrator with branch length proportional to the number of mutations specific to one tumor (private), two or more tumors (shared) or all tumors (trunk). Driver mutations and focal CNAs were added to the branches. All non-synonymous and synonymous mutations were used for tree construction. Signature analysis was performed for each individual tumor as well as trunks and each subsequent branch point for each tumor using deconstructSigs^75^. Mutations that were not signature 1, 2, 4, 5, or 13-type were labeled as “unclassified”. Mutational signature analysis was restricted to branches with at least 10 mutations. COSMIC mutational signatures were calculated for each branch using the R “deconstrucSigs” package^75^. Copy number phylogenetic trees were generated by hand. Branch lengths were drawn for visualization purposes only.

### Identification and classification of driver mutations

All identified nonsynonymous mutations were filtered to include only driver genes based on large-scale non-small cell lung cancer sequencing studies ^19,27,76–80^ and in the COSMIC cancer gene census (downloaded June 2016). We classified all nonsynonymous mutations into categories as previously described^1^. Category 1 ‘high-confidence driver mutations’ contained all disrupting mutations (nonsense, frameshift, splicing or ‘deleterious’ missense) in tumor suppressor genes or activating amino acid substitutions in non-small cell lung cancer oncogenes as described in lung cancer sequencing studies. Category 2 ‘putative driver mutations’ contained amino acid substitutions located at the same position or up to 5 amino acids away from a substitution present in COSMIC. Category 3 ‘low confidence driver mutations’ contained all other nonsilent mutations in genes that were present in the lists of cancer-related genes described above. Mutations were then analyzed using COSMIC to determine whether the amino acid substitution has been previously identified. Category 2 were further scored as ‘deleterious’ when at least two out of the three predictors classified the mutation as deleterious Functional prediction scores (SIFT, Polyphen2, and Provean). All category 1 mutations were considered deleterious and category 3 mutations were not included as driver mutations in any analyses.

### APOBEC germline allele determination

Germline *APOBEC3AB* deletion was genotyped by a proxy SNP rs12628403 and was genotyped using a custom-designed TaqMan genotyping assay, as described previously^30^. For patient RA006, the deletion status was also confirmed in all six tumors by Sanger sequencing, and expression analysis.

### qRT–PCR analysis

Total RNA for all experiments was isolated with the Qiagen All Prep DNA/RNA Mini Kit with on-column DNase I treatment. RNA quantity and quality as determined by RNA Integrity Number (RIN) were evaluated by Bioanalyzer RNA kit (Agilent). cDNA was prepared from equal amounts of total RNA for each sample with the RT2 first-strand cDNA kit and random hexamers with an additional DNA removal step (Qiagen). Expression of *APOBEC3A, APOBEC3B*, and *APOBEC3AB* deletion and endogenous controls *GAPDH* and *PPIA* was measured in each cDNA with TaqMan expression assays from Thermo Fisher: custom assays were used for *APOBEC3B:* (F: TGCTGGGAAAACTTTGTGTACAAT; R: ATGTGTCTGGATCCATCAGGTATCT; Probe: ATTCATGCCTTGGTACAAA), and *APOBEC3AB* (F: ATCATGACCTACGATGAATTTAAGCA; R: AGCACATTGCTTTGCTGGTG; Probe: FAM- CATTCTCCAGAATCAGGG), and commercial assays Hs00377444_m1 for *APOBEC3A*, 4326317E for *GAPDH* and 4326316E for *PPIA* (Thermo Fisher Scientific). Reactions were performed in four technical replicates on QuantStudio 7 (Life Technologies) using TaqMan Gene Expression buffer (Life Technologies); water and genomic DNA were used as negative controls for all assays. Expression was measured by Ct values (PCR cycle at detection threshold). Expression of *APOBEC3A, APOBEC3B* and *APOBEC3AB* was individually normalized by the mean of endogenous controls *(GAPDH* and *PPIA)*. Changes in expression were calculated using relative quantification method, as ΔCt = Ct (control) – Ct (target).

### Analysis of APOBEC mutagenesis

APOBEC-signature mutation analysis for all autopsy tumor samples was determined using an R software package kindly provided by Dr. Dmitry A. Gordenin^21,25,81^. We used two variables in the file *_sorted_sum_all_fisher_Pcorr.txt: the ‘tCw_to_G+tCw_to_T’ variable, which represents total counts of APOBEC-signature mutations, and the ‘APOBEC_Enrich’ variable, which accounts for statistical significance of enrichment and represents the level APOBEC mutagenesis pattern per sample. This second variable is more stringent, as many samples were not enriched at a statistically significant level and were classified as negative for APOBEC-signature mutations. We identified APOBEC ‘hypermutators’ as those with signature 2 + 13 mutations (TCGA and Broad datasets) or total APOBEC mutations (TRACERx and our dataset) exceeding 1.5 times the length of the interquartile range from the 75th percentile. This method to identify outliers has been previously described^22^.

We also used the same R software package to determine RT<u>C</u>A and YT<u>C</u>A enrichment for patient RA006. A less stringent filtering of whole-exome variants was used to provide sufficient sample size for this analysis. A Benjamin-Hochberg P value of 0.05 was used as a threshold for significance, unless specified otherwise, and all tests were two-sided.

### Copy Number Alteration Analysis

Copy number alteration (CNA) analysis was performed using MIP array technology (Affymetrix OncoScan FFPE Express 2.0) with 334,183 sequence tag site probes which were used to measure DNA copy number at different loci across the human genome. Copy number data were processed and normalized using the Affymetrix OSCHP-SNP-FASST2 algorithm within the Nexus Copy Number Software. We used a log2 ratio cut-off of +/− 0.5 to define focal copy number amplifications and deletions and +/− 0.25 to define arm-level copy number amplifications and deletions. To minimize overcalling heterogeneity of copy number alterations, we employed the following methods: 1) tumors without +/− 0.5 focal amplification/deletion were included if they had at log2 ratio +/− 0.2 and tumors without +/− 0.25 arm-level amplification/deletion were included if they had a log2 ratio +/− 0.10; 2) at least two tumors within a patient were required to have an amplification or deletion above the threshold of +/− 0.5 for focal and +/− 0.25 for arm; 3) an amplification/deletion was considered truncal if present in >80% of the tumors within a given patient. Allele-specific focal copy number profiles were determined for primary tumors (available for three patients) using the Sequenza package. Circos plots were generated using segmented GISTIC-output file for all tumors using circos v0.69–4, for every track the min and max are set to −1 and 1 respectively, values between −0.2 and 0.2 are not shown in the figure. Arm-level changes as depicted in the copy number phylogenetic trees were determined using GISTIC.

### RNA-sequencing and data processing

RNA-seq sequencing was performed on 31 out of 37 tumor sites. RNA-seq was done on Illumina HiSeq2500 platform to yield at least 100 million reads/sample using Illumina TruSeq V4 chemistry at 2 × 125 nt paired-end. Sequencing reads were aligned with TopHat version 2.0.13^82^ against the reference human genome hg19, with UCSC known gene transcripts as the gene model annotation. Expression on gene and isoform level was quantified with Cufflinks version 2.2.1^83^.

### RNA-seq variant calling and mutation validation

For RNASeq variants calling, sequencing reads were first aligned to hg19 with STAR version 2.4.2a and then with a second pass alignment to the transcriptome generated by STAR for each patient. For each identified SNV in WES, its expression was confirmed by the presence of sequencing reads of the alternative allele assessed by Samtools mpileup on TopHat generated BAM files from RNASeq data, whereas alternative reads coverage for indels were extracted from vcf files generated by the GATK best practices variant calling on RNASeq (see URLs). 69% of whole exome variants had a minimum 1X RNA depth and of these expressed variants 59% were confirmed by RNA-seq (Supplementary Fig. 2). 55% of whole exome variants had minimum of 5X RNA depth and of these expressed variants 69% were confirmed by RNA-seq (Supplementary Fig. 2). Validation rates for different variant types across all tumor samples were similar (range 42% silent to 56% nonsense) (Supplementary Fig. 2).

### RNA-seq data analysis

Cufflinks outputted FPKM values for each gene were normalized for all samples within each patient using limma package voom quantile method^84^. This expression data was used to predict enrichment scores among immune genes obtained from CIBERSORT for each sample^33^ and then using single-sample GSEA (ssGSEA) from GenePattern. Using the R package “fgsea”, GSEA preranked we performed to determine enrichment scores for REACTOME pathways. Principal component analysis (PCA) was used to combine clustered samples prior to conducting this analysis.

### Protein Extraction

All but one tumor (RA004 – Li1a) had sufficient tissue for mass-spectrometry (MS)-based proteomic characterization. About 10–15 mg of tumor tissue fresh-frozen in liquid nitrogen was lysed in 400μl of urea lysis buffer (20 mM HEPES pH 8.0, 8 M urea, 1 mM sodium orthovanadate, 2.5 mM sodium pyrophosphate and 1 mM ß-glycerophosphate) using a tissue lyser (Qiagen). Lysates were centrifuged at 14,000 rpm at 4ΰC for 10 mins and the clear supernatants were transferred to new tubes. Protein concentrations were determined by the Modified Lowry method (BioRad).

### Enzymatic Digestion

The protein lysate was reduced with 45 mM dithriothreitol (Sigma Aldrich, MO), alkylated with 100 mM iodoacetamide (Sigma Aldrich, MO), and subsequently digested with modified sequencing grade Trypsin (Promega, Madison, WI) at 30°C overnight. The digest was then acidified using 0.1% TFA and the peptides were desalted using solid phase extraction C18 column (Supelco, Bellefonte, PA), and vacuum dried in a centrifugal evaporator.

### TMT-Labeling

TMT10plex amine reactive reagents (0.8 mg per vial) (Thermo Fisher Scientific) were resuspended in 41 μL of anhydrous acetonitrile (ACN) and all 41 μL of each reagent was added to each sample and mixed briefly on a vortexer. Reactions were incubated at room temperature for 1 h, and then quenched by the addition of 8 μL of 5% hydroxylamine for 15 min and then combined at equal amount. All tumor tissues of LUAD patients RA000, RA003, RA005 and RA006 were pooled together to make a reference channel and labeled with TMT^10^–126. In a separate TMT labeling experiment, tumor tissues from patient RA006 were pooled together to make a reference channel.

### Basic reversed phase liquid chromatography (RPLC) fractionation

Basic RPLC separation was performed with a XBridge C18, 100 × 2.1 mm analytical column containing 5μm particles and equipped with a 10 × 2.1 mm guard column (Waters, Milford, MA) with a flow rate of 0.25 mL/min. The solvent consisted of 10 mM triethylammonium bicarbonate (TEABC) as mobile phase A, and 10 mM TEABC in ACN as mobile phase B. Sample separation was accomplished using the following linear gradient: from 0 to 1% B in 5min, from 1 to 10% B in 5min, from 10 to 35% B in 30min, and from 35 to 100% B in 5min, and held at 100% B for an additional 3min. A total of 96 fractions were collected during the LC separation in a 96-well plate in the presence of 12.5 μL of 1% formic acid. The collected fractions were concatenated into 12 fractions and dried in a vacuum centrifuge. One tenth of the peptides were injected directly for LC-MS/MS analysis.

### LC-MS/MS analyses

Peptides separated/fractionated by basic reversed-phase chromatography were analyzed on an LTQ-Orbitrap Elite interfaced with an Ultimate™ 3000 RSLCnano System (Thermo Scientific, San Jose, CA). The dried peptides were loaded onto a nano-trap column (Acclaim PepMap100 Nano Trap Column, C18, 5 μm, 100 Å, 100 μm i.d. × 2 cm) and separated on an Easy-spray^TM^ C18 LC column (Acclaim PepMap100, C18, 2 μm, 100 Å, 75 μm i.d. × 25 cm). Mobile phases A and B consisted of 0.1% formic acid in water and 0.1% formic acid in 90% ACN, respectively. Peptides were eluted from the column at 300 nL/min using the following linear gradient: from 4 to 35% B in 60min, from 35 to 45% B in 5min, from 45 to 90% B in 5min, and held at 90% B for an additional 5min. The heated capillary temperature and spray voltage were 275°C and 2kV, respectively. Full spectra were collected from m/z 350 to 1800 in the Orbitrap analyzer at a resolution of 120,000, followed by data-dependent HCD MS/MS scans of the fifteen most abundant ions at a resolution of 30,000, using 40% collision energy and dynamic exclusion time of 30s.

### Proteomic Data Analysis

Peptides and proteins were identified and quantified using the Maxquant software package (version 1.5.3.30) with the Andromeda search engine^85^. MS/MS spectra were searched against the Uniprot human protein database (May 2013, 38523 entries) and quantification was performed using default parameters for TMT10plex in MaxQuant. Corrected intensities of the reporter ions from TMT labels were obtained from the MaxQuant search. The relative ratios were calculated for each channel to the reference channel. These ratios were then used to predict enrichment scores of overall immune signatures obtained from CIBERSORT using single-sample GSEA (ssGSEA) from GenePattern and for REACTOME pathways by GSEA preranked through the R package “fgsea”. Samples were similarly combined as described for the RNA-seq data analysis. Cufflinks outputted FPKM values for each gene were normalized for all samples within each patient using limma package voom quantile method.

### Construction and Immunohistochemistry of Tissue Microarray

The physical construction of the TMA followed the guidelines previously used by the NCI Tissue Array Project. Each tumor from each autopsy patient was represented by 1 tumor core of 1mm that was taken from the original paraffin block. Serial 5μm sections were cut from the TMA block and used for immunohistochemical analysis. We used previously reported methods for immunohistochemical staining of TMAs^86^.

### Integrating copy number, gene expression and protein abundance

Pearson correlation coefficients (PCCs) were calculated across all common genes in copy number, gene expression and protein abundance data for each patient. Prior to calculating PCCs, gene expression data (RNA-seq FPKM) and protein abundance (protein ratios) were further normalized within each patient using the limma package with its voom quantile method. 3DPlots were created using the R package “scatterplot3d”. For arm-level analyses, normalized gene expression and protein abundance data was categorized by chromosomal arm. The mean of clusters of tumors, as determined previously by PCA, were calculated for both sets of data. Ratios were calculated between clusters and then log2 transformed. Probability density plots were generated with 1% outliers removed and x-axis of plots were restricted to −1 to 1 (log2 scale) for gene expression and −0.1 and +0.1 (log2 scale) for protein abundance for visualization purposes.

### Statistics and graphics

All figures and graphs were generated using the “ggplot2” package available through the R statistical program. Linear regression, correlations and t-tests were conducted though the R base packages. All tests were two-tailed and p-values less than 0.05 were considered significant.

### URLs

Firehose Broad GDAC, https://gdac.broadinstitute.org/

Webgestalt, http://webgestalt.org/

Gene pattern, https://software.broadinstitute.org/cancer/software/genepattern/

SnpEff, http://snpeff.sourceforge.net

Gatk, https://software.broadinstitute.org/gatk/

National Cancer Institute’s Tissue Array Project, http://ccr.cancer.gov/tech_initiatives/tarp/default.asp

### Data Availability

The sequencing and genotype data have been deposited at the database of Genotypes and Phenotypes (dbGaP), which is hosted by the National Center for Biotechnology Information (NCBI), under accession number phs001432.v1.p1.

## AUTHOR CONTRIBUTIONS

N.R. and U.G. designed the study. N.R., T.K.M., X.Z., A.V., C.M.C., A.R.B., L.P-O. and U.G. assisted with sample preparation and data analysis. N.R., T.K.M, X.Z. and U.G. performed the proteomic experiments and data analysis. N.R., S.G., R.P., S.S., A.G., A.R.P. and J.K. performed or assisted with computational analysis. N.R. performed the biostatistical analyses. S.H. and D.K. provided guidance and assisted with autopsies. A.V, T.K.M, R.B and U.G. harvested tissues during autopsies. A.T., A.R., C.A.C and U.G. were involved with clinical care of patients undergoing autopsies. N.R. and U.G. wrote the manuscript. All authors reviewed, commented on, and approved the manuscript.

## ACKNOWLEDGMENTS

We thank Susan Perry, Emerson Padiernos, the clinical and research nurses as well as palliative care and spiritual care teams for their clinical care of the patients in this study who were enrolled in hospice prior to autopsy. We also thank Willie Young and the Pathology residents who assisted with the autopsies. We thank Dr. Dmitry A. Gordenin, Dr. Kin Chan, Dr. Bing Zhang and Dr. Jing Wing for their helpful discussions.

## Supplementary Table Legends

Supplementary Table 1: Clinical characteristics, autopsy findings and treatment history of all study patients

Supplementary Table 2: Variants identified within each patient by whole-exome and RNA-sequencing

Supplementary Table 3: Jaccard similarity coefficients of metastases within each patient based on whole-exome, deep whole-exome and RNA sequencing

Supplementary Table 4: Summary of APOBEC mutagenesis within each tumor based on whole-exome and RNA-sequencing

Supplementary Table 5: Log2 copy number ratios of chromosomal arms within each patient

Supplementary Table 6: Normalized FPKM of all genes within each patient categorized by chromosomal arm location

Supplementary Table 7: Normalized protein abundance data of all genes within each patient categorized by chromosomal arm location

Supplementary Table 8: Log2 copy number ratios of amplified or deleted genes previously determined to be significant in lung cancer by GISTIC

Supplementary Table 9: Common pathways identified by gene set enrichment analysis within transcriptomic and proteomic datasets between tumors of each patient

